# Decoding murine cytomegalovirus

**DOI:** 10.1101/2022.11.10.515907

**Authors:** Manivel Lodha, Ihsan Muchsin, Christopher Jürges, Vanda Juranic Lisnic, Anne L’hernault, Andrzej J. Rutkowski, Bhupesh Prusty, Arnhild Grothey, Andrea Milic, Thomas Hennig, Stipan Jonjic, Caroline C. Friedel, Florian Erhard, Lars Dölken

## Abstract

The genomes of both human cytomegalovirus (HCMV) and murine cytomegalovirus (MCMV) were first sequenced over 20 years ago. Similar to HCMV, the MCMV genome had initially been proposed to harbor ≈170 open reading frames (ORFs). More recently, omics approaches revealed HCMV gene expression to be substantially more complex comprising several hundreds of translated ORFs. Here, we provide a state-of-the art reannotation of lytic MCMV gene expression based on integrative analysis of a large set of omics data. Our data reveal 363 viral transcription start sites (TiSS) that give rise to 380 and 454 viral transcripts and ORFs, respectively. The latter include >200 small ORFs, some of which represented the most highly expressed viral gene products. By combining TiSS profiling with metabolic RNA labelling and chemical nucleotide conversion sequencing (dSLAM-seq), we provide a detailed picture of the expression kinetics of viral transcription. This not only resulted in the identification of a novel MCMV immediate early transcript encoding the m166.5 ORF, which we termed *ie4*, but also revealed a group of well-expressed viral transcripts that are induced later than canonical true late genes and contain an initiator element (Inr) but no TATA- or TATT-box in their core promoters. We show that viral uORFs tune gene expression of longer viral ORFs expressed in *cis* at translational level. Finally, we identify a truncated isoform of the viral NK-cell immune evasin m145 arising from a viral TiSS downstream of the canonical m145 mRNA. Despite being ≈5-fold more abundantly expressed than the canonical m145 protein it was not required for downregulating the NK cell ligand, MULT-I. In summary, our work will pave the way for future mechanistic studies on previously unknown cytomegalovirus gene products in an important virus animal model.

**Author summary:** We conducted a comprehensive characterization and reannotation of murine cytomegalovirus (MCMV) gene expression during lytic infection in murine fibroblasts using an integrative multi-omics approach. This unveiled hundreds of novel transcripts that explained the expression of close to 300 so far unknown viral open reading frames (ORFs). Interestingly, small viral ORFs (sORFs) were amongst the most highly expressed viral gene products and thus presumably encode for important viral microproteins of unknown function. However, we also show that sORFs located upstream of larger ORFs tune the expression of the downstream ORFs at the level of translation. We classified viral transcription start sites (TiSS) based on their expression kinetics obtained by a new combination of metabolic RNA labelling with transcription start sites profiling. This not only identified a so far unknown viral immediate-early transcript (*ie4*, m166.5 RNA) but also revealed a novel class of viral late transcripts that are expressed later than canonical true late genes and lack TATA box-like motifs. We exemplify for the m145 locus how so far unknown TiSS give rise to abundantly expressed truncated viral proteins. In summary, we provide a state-of-the-art annotation of an important model virus, which will be instrumental for future studies on CMV biology, immunology and pathogenesis.

## Introduction

Human cytomegalovirus (HCMV) is a ubiquitous pathogen that establishes a life-long infection upon primary infection [1]. While primary infection is mostly asymptomatic, HCMV is responsible for a significant morbidity and mortality in immunocompromised patients and neonates. There is currently no vaccine. The strict species specificity of HCMV poses a major challenge in understanding cytomegalovirus (CMV) pathogenesis [2]. Murine cytomegalovirus (MCMV) exhibits significant similarity to HCMV and represents a widely used model to study CMV pathogenesis [2, 3]. CMV gene expression is temporally regulated and classified into immediate early (IE), early (E) and late (L) gene expression [4]. In contrast to viral *ie* gene expression, the expression of E genes requires *de novo* expression of the major viral transcription factor IE3 and thus viral protein synthesis [5]. Viral L gene expression depends on viral DNA replication as well as expression of the viral late gene transcription factor (LTF) complex that binds to a TATA-like (TATT) motif in the proximal promoters of viral late genes [6–10].

In recent years, high-throughput sequencing technologies, including ribosome profiling (Ribo-seq) [11] and RNA-seq [12] reshaped our understanding of the coding capacity of herpesviruses including HCMV [13], HSV-1 [14], KSHV [15] and EBV [16]. Strikingly, these studies revealed the presence of hundreds of novel open reading frames (ORFs). These predominantly arise from promiscuous transcription initiation within the viral genome. Many of these novel ORFs are small ORFs (sORFs) of <100 amino acids (aa) in size. Depending on their genome location with respect to the larger viral ORFs, they are referred to as upstream ORFs (uORFs), upstream overlapping ORFs (uoORFs), internal ORFs (iORFs), or downstream ORFs (dORFs) [14, 17].

The 230-kb genome of the MCMV Smith strain was initially predicted to encode 170 protein coding sequences (CDS), many of which share homology to HCMV [18]. To date, a state-of-the art reannotation of the MCMV genome including mRNAs, short ORFs and isoforms of canonical ORFs as well as an overarching hierarchical nomenclature has been lacking. Nevertheless, additional viral gene products have been discovered through various genetic [19, 20], *in silico* [21] and proteomic approaches [22]. This also includes the identification of different MCMV protein isoforms, which arise from alternative viral transcripts expressed with distinct kinetics [23]. A prominent example for the need for comprehensive annotation of the MCMV genome was the identification of the 83-amino acid microprotein MATp1 [24]. MATp1 is expressed from the most abundant MCMV transcript (MAT) upstream of the coding sequence (CDS) of the spliced m169 gene [25]. Initially dismissed by *in silico* predictions due to its small size (≈83 aa), MATp1 acts in concert with the viral m04 protein and specific MHC-I allotypes in a trimeric complex to evade missing-self recognition by natural killer (NK) cells [24]. Furthermore, recognition of this trimeric complex by at least three activating NK-cell receptors explains intrinsic resistance of certain mouse strains to MCMV infection [24]. These findings highlight the importance of studying gene expression at single nucleotide resolution using unbiased, integrative multi-omics approaches to fully understand the coding potential of MCMV. The wealth of novel viral gene products requires a revised nomenclature.

We recently utilized a multi-omics approach coupled with integrative computational analysis to decipher the transcriptome and translatome of herpes simplex virus 1 (HSV-1) [14]. Here, we use a similar approach to comprehensively identify, characterize and hierarchically annotate MCMV gene products expressed during lytic infection of murine NIH-3T3 fibroblasts **(Fig. 1).** Our new annotation comprises 363 viral transcription start sites (TiSS) that give rise to 380 and 454 viral transcripts and ORFs, respectively. TiSS profiling combined with metabolic RNA labelling and chemical nucleotide conversion sequencing (dSLAM-seq) resolved the kinetics of viral gene expression and their regulation by core promoter motifs. Abundant transcription initiation and alternative TiSS usage throughout lytic infection explained the expression of hundreds of novel viral ORFs and small ORFs, as well as N-terminal extensions (NTE) and truncations (NTT) thereof revealed by ribosome profiling. In summary, our work provides a state-of-the-art annotation of an important virus model.

**Fig 1.**
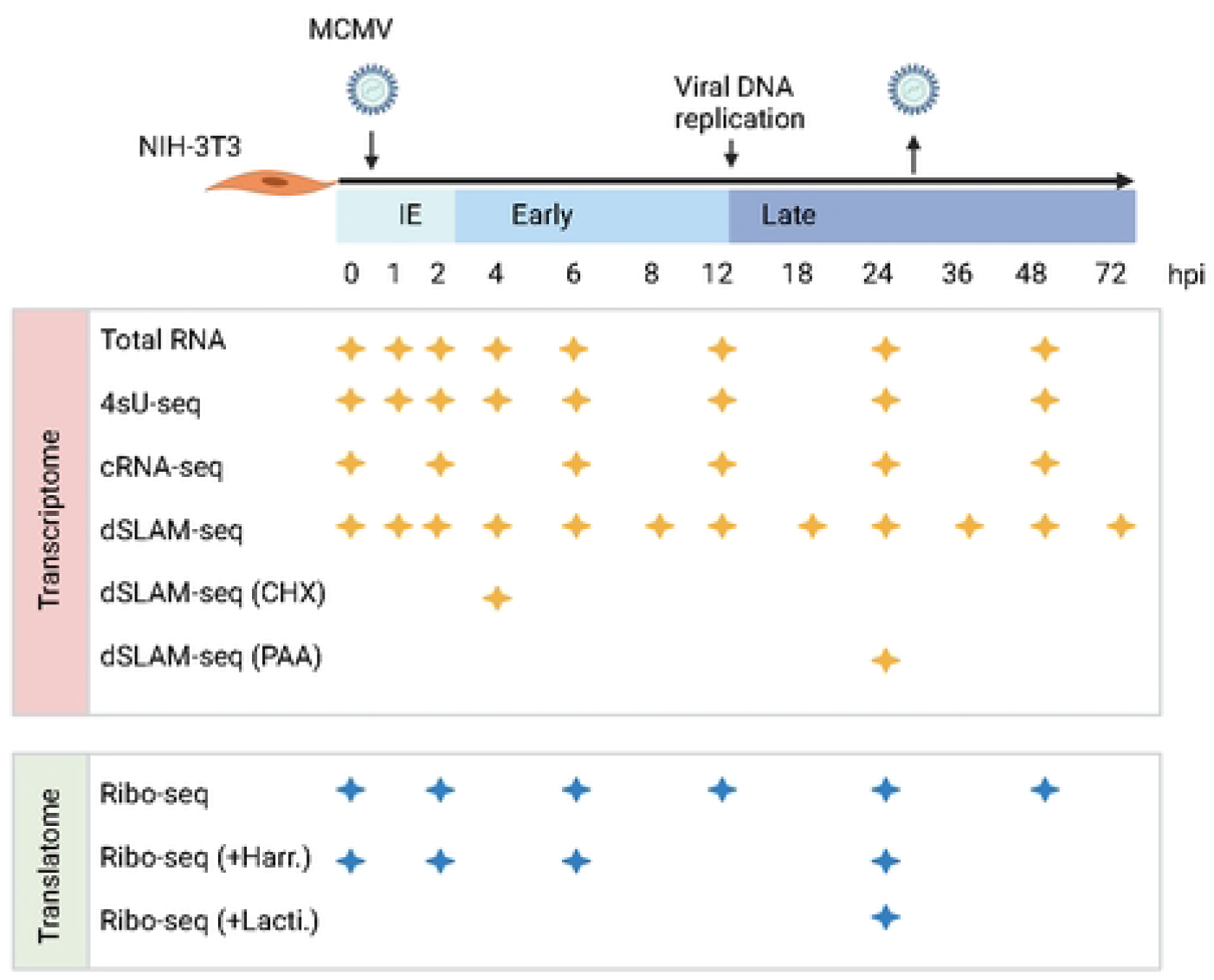
Overview of applied omics approaches. MCMV gene expression was analyzed in Swiss murine embryonic fibroblasts (NIH-3T3) infected with BAC-derived wild-type MCMV at an MOI of 10. Viral transcription start sites (TiSS) and splicing events were determined through total RNA-seq, 4sU-seq, cRNA-seq and dSLAM-seq (n=2; including one biological replicate for dSLAM-seq with cycloheximide (CHX; 4 h) or phosphonoacetic acid (PAA; 24 h) treatment). To decipher the MCMV translatome, four biological replicates of ribosome profiling were performed. Enrichment of reads at translation start sites (TaSS) was improved by pre-treating cells with Harringtonine – Harr. (two biological replicates) or Lactimidomycin – Lacti. (one biological replicate) for 30 min. The available time points and conditions are indicated by stars for any given approach.

## Results

### Characterization of the MCMV transcriptome

To identify the full complement of MCMV transcripts in lytic infection of fibroblasts, we profiled viral gene expression in MCMV-infected NIH-3T3 fibroblasts throughout the first three days of infection using multiple next-generation sequencing approaches **(Fig. 1)**. This included: (***i***) RNA-seq of total RNA (Total RNA-seq) and (***ii***) newly transcribed RNA obtained by metabolic RNA labelling using 4-thiouridine (4sU-seq) [26] from the same samples. To analyze temporally resolved promoter usage, we performed transcription start sites (TiSS) profiling by (***iii***) cRNA-seq [13, 14] as well as (***iv***) dSLAM-seq, a novel combination of differential RNA-seq (dRNA-seq) [27] with metabolic RNA labelling and thiol(SH)-linked alkylation of RNA (SLAM-seq) [28]. A representative example of the obtained data are shown for the M25 locus in **Fig. 2A**. cRNA-seq is a modified total RNA sequencing protocol that is based on circularization of RNA fragments (hence termed cRNA-seq) [14]. It allows both TiSS identification based on a moderate enrichment (median: 8-fold) of reads starting at 5’ RNA ends (**Fig. 2B)** and quantification of total transcript levels. In contrast, dSLAM-seq provides a much greater enrichment of TiSS (median: 24-fold) by selectively enriching reads at 5’ ends of cap-protected RNA fragments resistant to 5’-3’ Xrn1 exonuclease digest (**Fig. 2B**) **(S1A Fig.)**. Importantly, dSLAM-seq combines 1 h 4sU labelling immediately prior to cell lysis, followed by RNA isolation and chemical conversion of the introduced 4sU residues to a cytosine analogue (SLAM-seq). The latter facilitates computational identification of sequencing reads derived from newly transcribed RNA (new RNA) based on the introduced U-to-C conversions [29]. Selective analysis of new RNA in dSLAM-seq data thus reveals the true temporal kinetics of gene expression for each viral TiSS. In addition, we also included dSLAM-seq samples pre-treated with chemical inhibitors of protein synthesis and viral DNA replication, namely cycloheximide (CHX; 4 hours post infection (hpi)) and phosphonoacetic acid (PAA; 24 hpi), respectively. A detailed overview of the analyzed time points and conditions is shown in **Fig. 1.**

**Fig 2.**
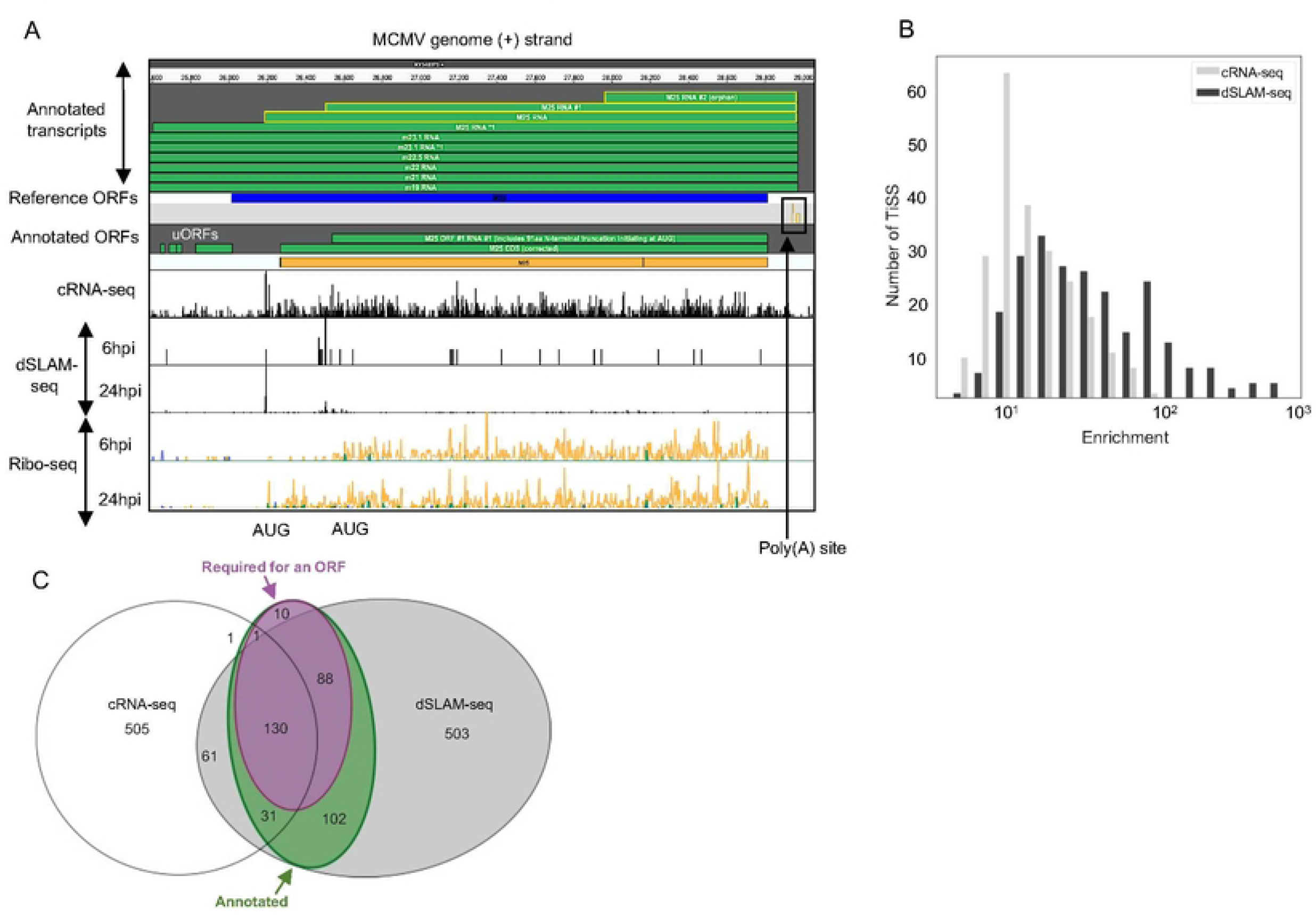
Characterization of the MCMV transcriptome. **A.** Screenshot of MCMV gene expression showing annotated transcripts and ORFs in the M25 locus with 5’ read enrichment at TiSS as depicted by cRNA-seq and dSLAM-seq as well as Ribo-seq data, respectively. Viral transcripts that initiate within the depicted region of the MCMV genome are highlighted in yellow. The schematic portrays translation of the 130 and 105 kDa M25 protein isoforms validated in a recent study [23]. The M25 RNA *1 also encodes four small ORFs (M25 uORFs 1-3 and M25 uoORF) of 6, 11, 8 and 63 aa, respectively. dSLAM-seq data are depicted in linear scale, Ribo-seq data in logarithmic scale. **B.** Graphical representation of 5’ read enrichment obtained by dSLAM-seq and cRNA-seq approaches. **C.** Venn diagram depicting the number of TiSS identified by both cRNA-seq and dSLAM-seq. TiSS included in the final annotation are depicted in the green circle as “annotated”. TiSS labelled as “Required for an ORF” represent TiSS that are required to explain the translation of a downstream ORF (no other TiSS within 500 nt upstream of the ORF).

Reliable identification of viral TiSS requires integrative analysis of multiple data sets from different TiSS profiling approaches and kinetic studies [14]. We thus employed our recently published integrative TiSS analysis pipeline iTiSS [30], which identifies statistically significant peaks arising from TiSS profiling read accumulations on the genome. We furthermore scored these TiSS candidates according to a variety of additional criteria, including an increase in upstream to downstream read coverage in cRNA-seq and 4sU-seq data, temporal changes in cRNA-seq and dSLAM-seq read counts and the presence of translated ORFs identified by Ribo-seq, for which no other transcript could otherwise be identified. This resulted in a maximum score of 7 for any given candidate TiSS. We then manually inspected all candidate TiSS using our in-house MCMV genome browser, which combines all data sets, time points and conditions (**Fig. 2A**). In total, we identified and annotated 363 unique MCMV TiSS **(Fig. 2C**), satisfying the given set of criteria **(S1B Fig.)**. Some TiSS were common for alternatively spliced products and differential poly(A) site usage resulting in a total of 380 MCMV transcripts. The complete list of all MCMV transcripts and their respective scores is included in **S1 Table**.

We next analyzed splicing events in the MCMV transcriptome based on our total RNA-seq and 4sU-seq data. We first identified all unique reads spanning exon-exon junctions by at least 10 nt (see **Methods** for details). We identified 366 splicing events, most of which only occurred at very low levels **(Fig. 3)** and had no impact on the identified ORFs. We thus decided not to include them into our new reference annotation and only retained 27 splicing events. Six of these splicing events had already been reported by Lisnic *et al*. [25] and several of these had been successfully validated using RT-PCR and 3’ sequencing in the same study as well as other studies **(S2 Table)**. To independently validate the identified splice sites and investigate the impact of the corresponding transcript isoforms on translation, we utilized our Ribo-seq data. We confirmed an alternative splice donor site in the m133 locus as suggested by Rawlinson *et al*; [18] leading to the expression of two protein isoforms from differentially spliced ORFs **(S2A Fig.)**. Splicing of both the most abundant transcript (MAT) within the m169 locus and of a highly expressed transcript in the M116 locus were readily confirmed in our data **(S2B Fig.)**, the latter readily explained a recently validated spliced protein, M116.1p, which was found to be crucial for efficient infection of mononuclear phagocytes [31]. We also confirmed a previously reported spliced ORF in the m147.5 locus [32] **(S2C Fig.)** along with a novel splicing event in the m124 locus, leading to a correction of the previously annotated m124 ORF [18] **(S2D Fig.)**. While we readily observed the well-described MCMV 7.2 kb intron [33, 34], we were unable to detect the overlapping 8 kb intron reported in the same study [33]. 4sU-seq data also revealed multiple alternative donor sites in the m60-m73.5 locus **(S3 Fig.)**, which expressed several weakly expressed isoforms of the m73.5 ORF, the most dominant being the M73-m73.5 spliced ORF. Our data demonstrate that splicing in the MCMV transcriptome is much more prevalent than previously thought but mostly comprises low level splicing events in addition to the previously described splicing events. A complete list of annotated splicing events, which we included into our new reference annotation of the MCMV genome, is included in **S2 Table** and a list of all 366 putative 4sU-seq based introns are included in **S3 Table**.

**Fig 3.**
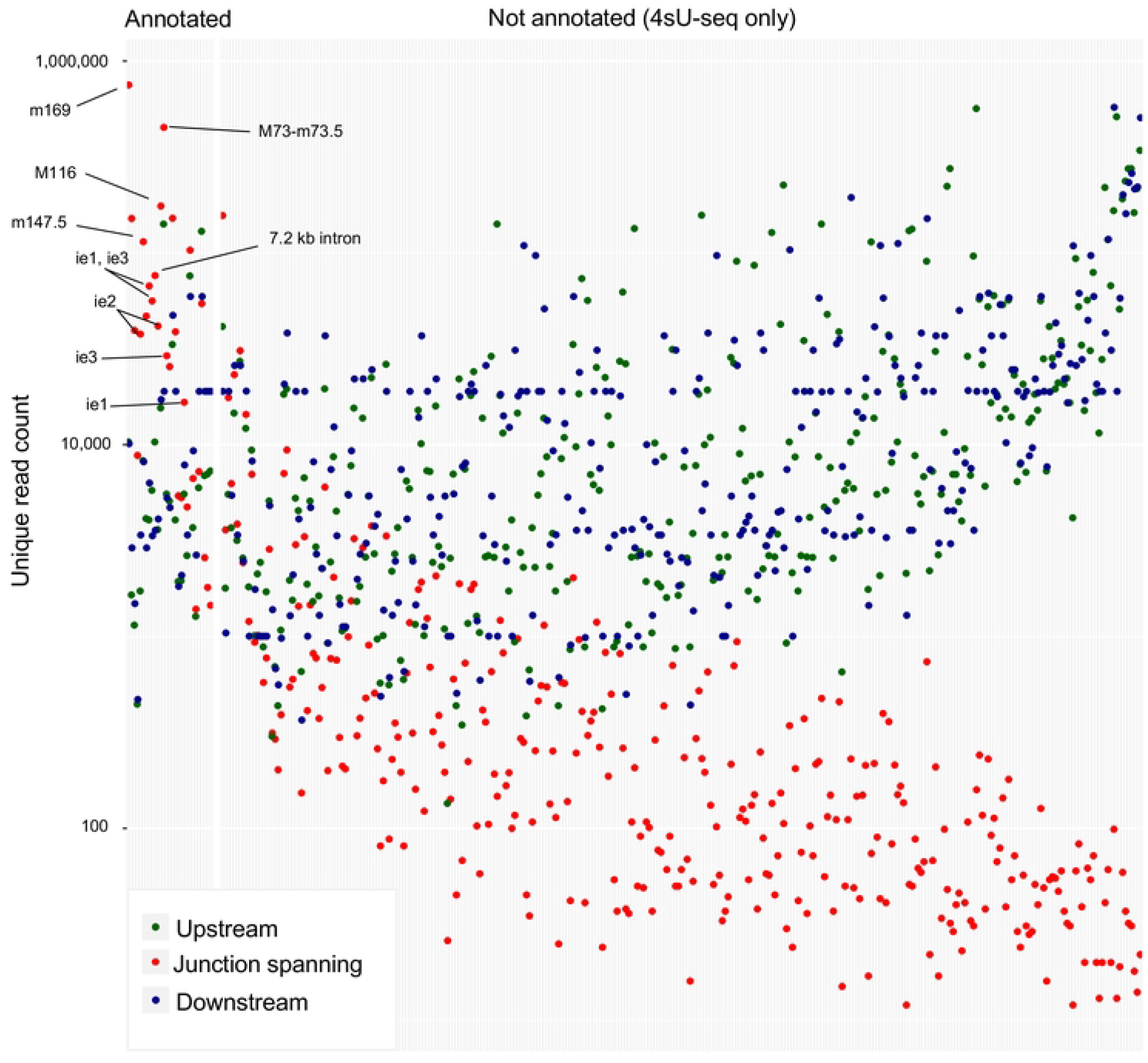
Identification of MCMV splicing events. Mapped reads from 4sU-seq and total RNA-seq identified 366 putative splicing events in the MCMV transcriptome. The y-axis displays the number of reads occurring at a spliced region, further categorized into reads spanning exon-exon junctions (red) by at least 10 nt as well as non-exon-spanning reads upstream (green) and downstream (blue). Putative splicing events were sorted based on the ratio of spliced (red) to unspliced (green + blue) reads. Only 27 of the 366 putative splicing events were included into our new reference annotation because they (***i***) had already been identified by others (16/27; **S2 Table**), (***ii***) were highly abundant, or (***iii***) affected the coding sequence of an MCMV ORF or sORF. To avoid unnecessary complexity in the revised annotation of the MCMV transcriptome, we excluded the other (putative) splicing events from our new reference annotation.

### Temporal regulation of viral transcription

Many core promoters of eukaryotic genes contain TATA boxes [35], which are also prevalent in herpesvirus genomes [14]. Utilizing total RNA levels obtained from our MCMV dSLAM-seq data, we grouped transcripts according to levels of gene expression (high, mid and low transcription). T/A rich regions indicative of TATA box-like motifs were much more prevalent in highly transcribed viral genes than in lowly transcribed genes **(Fig. 4A)**. In mammalian cells, TiSS are marked by an initiator element (Inr), characterized by a pyrimidine-purine dinucleotide [36]. As previously observed for HSV-1 [14], Inr elements were also prevalent for MCMV TiSS irrespective of their expression levels. This confirmed reliable identification of TiSS even for the most weakly utilized viral TiSS.

**Fig 4.**
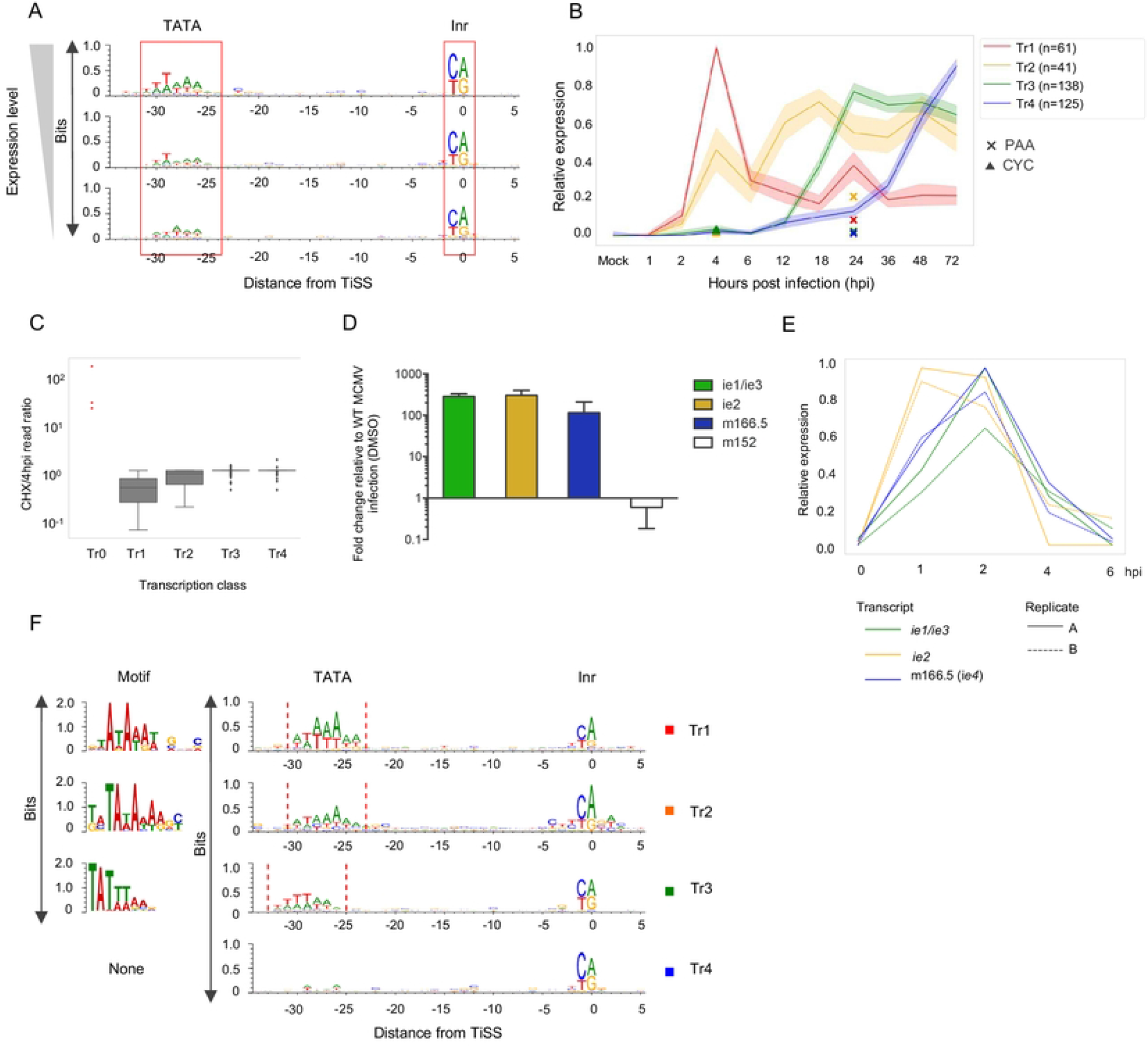
dSLAM-seq reveals distinct core promoter motifs associated with viral gene expression kinetics and a novel viral *ie* gene (*ie4*). **A.** Depiction of core promoter motifs of viral TiSS clustered according to their transcription rates in three equally sized bins (high, mid and low). The TATA box and initiator element (Inr) are shown. **B.** Graphical depiction of 4 clusters (Tr1-4) of viral TiSS obtained based on new RNA derived from the dSLAM-seq data. PAA (24 h) and CHX (4 h) treatments are indicated by a star and triangle, respectively. **C.** Ratio of new RNA levels with and without CHX pre-treatment were compared for various transcription classes (Tr0-Tr4). Tr0 genes are indicated as a dot-plot representing the *ie1/ie3*, ie2 and *ie4* (m166.5) TiSS. **D.** Validation of the m166.5 RNA as an *ie4* gene by qPCR. qPCR was performed with and w/o cycloheximide treatment on MCMV infected NIH-3T3 cells harvested 4hpi. GAPDH was used as a housekeeping gene and results were plotted as fold change relative to MCMV infection under DMSO treatment for three biological replicates. **E.** Line graphs representing relative levels of *ie* gene expression during early stages of infection. **F.** Graphical and sequence logo depiction of core promoter motifs identified for Tr1-4 clusters through MEME motif analysis. Please note that the TATA-/TATT-box motif in cluster Tr3 is shifted by 2 nt to the left compared to cluster Tr1.

Metabolic RNA labelling and chemical nucleotide conversion combined with dSLAM-seq enabled us to analyze real-time transcriptional activity of each individual viral TiSS throughout the course of lytic infection. We clustered viral TiSS according to their temporal regulation in “new RNA” and obtained four distinct clusters consistent with early (Tr1), early-late (Tr2), late (Tr3) and late* (Tr4) transcripts **(Fig. 4B)**.

MCMV immediate early genes (*ie1, ie2* and *ie3*) do not require viral protein synthesis and are thus resistant to inhibition of protein synthesis by cycloheximide (CHX). To identify novel MCMV immediate early genes, our dSLAM-seq experiment included a single replicate of 4 h of CHX treatment, which was initiated at the time of infection. Interestingly, CHX treatment not only confirmed the two immediate early TiSS of *ie1/ie3* and *ie2* but revealed transcription of one additional TiSS, namely the m166.5 RNA encoding for the m166.5 ORF of 446 aa (**Fig. 4C, S4A Fig.)**. qRT-PCR analysis confirmed these findings revealing a ~100-fold increased TiSS usage upon CHX treatment by 4 hpi compared to the untreated control (**Fig. 4D**). We thus termed m166.5 immediate early gene 4 (*ie4*). The respective m166.5 ORF (IE4) has been shown to encode a nuclear protein [19], but lacks functional characterization. In contrast to the other MCMV *ie* genes, *ie4* does not contain any introns. Interestingly, all three immediate early TiSS (*ie1/ie3, ie2* and ie4) show near identical kinetics throughout the full course of lytic MCMV infection. This included an early peak at 2 hpi and a low at 6 hpi (**Fig. 4E**) followed by a continuous rise until late in infection (72 hpi) (**S4B Fig.**). We thus defined the three immediate early TiSS as “Tr0” (**S4 Table**).

Tr1 expression peaked at 4 hpi followed by strong downregulation in transcriptional activity despite viral DNA replication. In contrast, Tr2 expression only showed a minor dip at 6 hpi and then reached a plateau by 12 hpi that remained remarkably stable until late in infection. In contrast, to CMV IE and E genes, viral L genes require viral DNA replication as well as the late viral transcription factor complex (LTF) [37]. The highly conserved CMV LTF is comprised of six viral proteins and binds to a modified TATA-box, i.e. a TAT<u>T</u> motif [10]. While canonical TATA boxes were a hallmark motif of Tr1 transcripts both TATA and TATT motifs were enriched for Tr2 transcripts. Clusters Tr1 and Tr2 thus comprise canonical MCMV early and early-late genes, respectively. In contrast, promoters of Tr3 transcripts harbored TATT motifs **(Fig. 4F)**. The Tr3 cluster comprises the canonical MCMV late genes, which commonly encode for structural proteins like the small capsid protein (SCP). Their expression is driven by the viral LTF complex. To assess the impact of TATA and TATT motifs on the kinetics and extent of viral gene expression, we utilized a dual color reporter virus (MCMV_Δm152-EGFP_SCP-IRES-mCherry). This virus expresses eGFP instead of the coding sequence of the m152 early gene (Tr1) and mCherry expressed from an internal-ribosomal entry site (IRES) downstream of the late gene m48.2 CDS encoding for SCP (Tr3). We mutated the TATA box of the m152 promoter to create a TATT motif. Upon infection of NIH-3T3 cells with the two viruses for 6 to 72 hours, we analyzed eGFP and mCherry fluorescence by microscopy (**Fig. 5A**) and flow cytometry (**Fig. 5B**). Consistent with our dSLAM-seq data, hardly any mCherry expression was observed within the first 12 h of infection and subsequent mCherry expression was sensitive to inhibition of viral DNA replication by PAA treatment. Interestingly, the TATA>TATT single point mutation was sufficient to render the m152 promoter PAA-sensitive. However, the TATA>TATT mutation only altered the kinetics but not the maximum mean fluorescence intensity (MFI) of eGFP expression throughout infection. Furthermore, introduction of the TATT-motif did not abrogate m152 promoter activity early in infection and only reduced total eGFP expression levels during the first 12 h of infection by ≈2-fold. Thus, while the TATT-motif defines sensitivity of a promoter to viral DNA replication, other promoter motifs or features define viral promoter activity during the early phase of infection.

**Fig 5.**
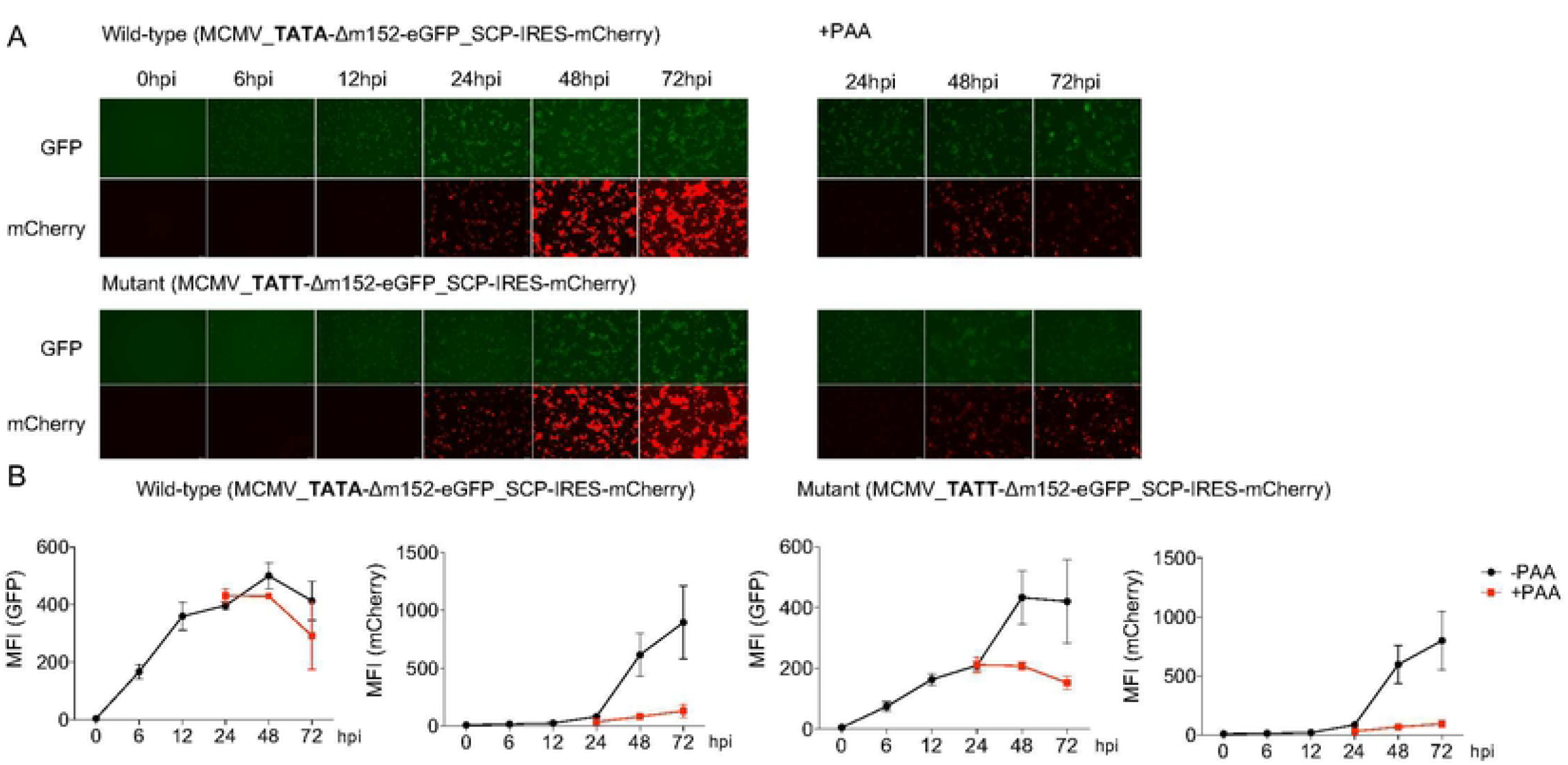
Converting a TATA box to a TATT box is sufficient to alter viral gene expression kinetics. **A.** NIH-3T3 cells were infected with a two-color MCMV reporter virus (MCMV_TATA-Δm152-eGFP_SCP-IRES-mCherry) and the TATA>TATT mutant thereof (MCMV_TATT-Δm152-eGFP_SCP-IRES-mCherry) viruses at an MOI of 1 for the indicated time points with and without PAA treatment. mCherry and eGFP expression was analyzed through fluorescence microscopy. Representative images of three biological replicates (n=3) are shown. **B.** Cells were fixed and eGFP and mCherry levels were analyzed quantitatively through flow cytometry and mean fluorescent intensity (MFI) values were plotted for three biological replicates (n=3) along with standard deviation (S.D.) with and without PAA pre-treatment.

Interestingly, the Tr4 cluster was characterized by late onset and the absence of TATA or TATT motifs. The latter is indicative of LTF-independent transcription at late stages of infection. Nevertheless, transcription of the Tr4 cluster was significantly impaired upon inhibition of viral DNA replication by PAA **(Fig. 4B)**. Interestingly, expression of Tr4 transcripts increased significantly later than of Tr3. However, Tr4 transcripts were also expressed at substantially lower levels than transcripts in Tr3 (**S5A, B Fig.**), possibly due to the absence of TATA or TATT motifs. This raised the question whether Tr4 transcripts are indeed regulated differently than Tr3 transcripts or whether their distinct kinetics are only observed due to lower transcriptional activity. To discern Tr4 as an independent cluster, we segregated transcripts in the Tr3 and Tr4 clusters into four quartiles according to levels of TiSS expression using our dSLAM-seq data. Tr3 and Tr4 transcripts exhibited distinct kinetics for all quartiles (**S5A Fig.**). Furthermore, even the least strongly expressed genes in Tr3 were associated with a distinctly positioned TATT motif, while even the most highly expressed Tr4 transcripts were not (**S5B Fig.**). Finally, the top thirty transcripts of Tr4 showed no evidence of a TATT motif, while 30 expression-matched Tr3 transcripts were still associated with a prominent TATT motif **(S5C Fig.)**. Our data thus reveal a new class of MCMV late transcripts that lack a distinct TATA/TATT and are expressed with delayed kinetics compared to the canonical late transcripts. We thus decided to term Tr4 as “late*” transcripts. It is important to note that the classical MCMV late transcripts that encode for canonical structural virion proteins belong to Tr3 and not Tr4. We hypothesize that transcription initiation of Tr4 transcripts does not require a TATT motif but is predominantly driven by the excessive amounts of viral DNA at late stages of infection. It will be interesting to assess (***i***) the relevance of viral Tr4 gene expression for productive infection and (***ii***) whether it is indeed independent of the viral LTF.

### Decoding the MCMV translatome

To decode the MCMV translatome, we employed ribosome profiling (Ribo-seq) along with translation start site (TaSS) profiling [38] across a time course of MCMV-infected NIH-3T3 cells **(Fig. 1)**. We identified and annotated a total of 454 MCMV ORFs including 227 small ORFs **(S5 Table)**. Using the annotation described by Rawlinson *et al*. [18] as reference, we confirmed 150 out of the 170 predicted CDS **(Fig. 6A)**. Putative CDS with no signs of translation are included in **S6 Table**. Interestingly, most of the predicted CDS that we were unable to detect were low-scoring predictions as per previously described criteria and no corresponding TiSS could be identified. The absence of corresponding transcripts in MCMV infection of fibroblasts explains the absence of detectable levels of translation. As the respective transcripts may be expressed in other cell types or conditions, we nevertheless maintained these CDS in our new genome annotation but labelled them as “orphan; not expressed”. Additionally, we detected 11 previously validated ORFs **(S7 Table)**. Overall, we identified 50 novel large ORFs, 10 N-terminal extensions (NTE), 27 N-terminal truncations (NTT) and 227 small ORFs of <100 aa (including upstream ORFs (uORFs), upstream overlapping ORFs (uoORFs), internal ORFs (iORFs), downstream ORFs (dORFs) and other short ORFs (sORFs) **(Fig. 6B)**. Specific viral transcripts initiating less than 500 nt upstream of the respective ORFs explained translation of 366 of 454 MCMV ORFs. Only for 88 viral ORFs, (66 of 232 small ORFs), for which no TiSS could be identified within the upstream 500 nt, were also included as “orphans” in our final annotation. The majority (50 of 57, 88%) of large viral ORFs initiated at canonical AUG start codons. Alternative start codons included ACG (1 ORFs/9 sORFs), GUG (1 ORFs/ 4 sORFs) and CUG (5 ORFs/ 10 small ORFs), with 12% of novel large ORFs **(Fig. 6C)** and 12 % of novel small ORFs **(Fig. 6D)** initiating at non-canonical codons. Most of the 27 NTTs and 10 NTEs identified by our pipeline resulted from alternative TiSS usage. Consistent with the rules applied for CDS identification by Rawlinson et al. [18], all NTTs initiated from AUG start codons whereas NTEs predominantly initiated at non-canonical start codons, which had previously not been considered **(Fig. 6E, F)**. We identified an N-terminal truncation in the *ie2* locus, i.e., m128 CDS #1 RNA #1 initiating at a novel early transcript (m128 RNA #1), which confirms previous observations of a modified IE2 protein of 41 kDa [39]. **(S6 Fig.)**

**Fig 6.**
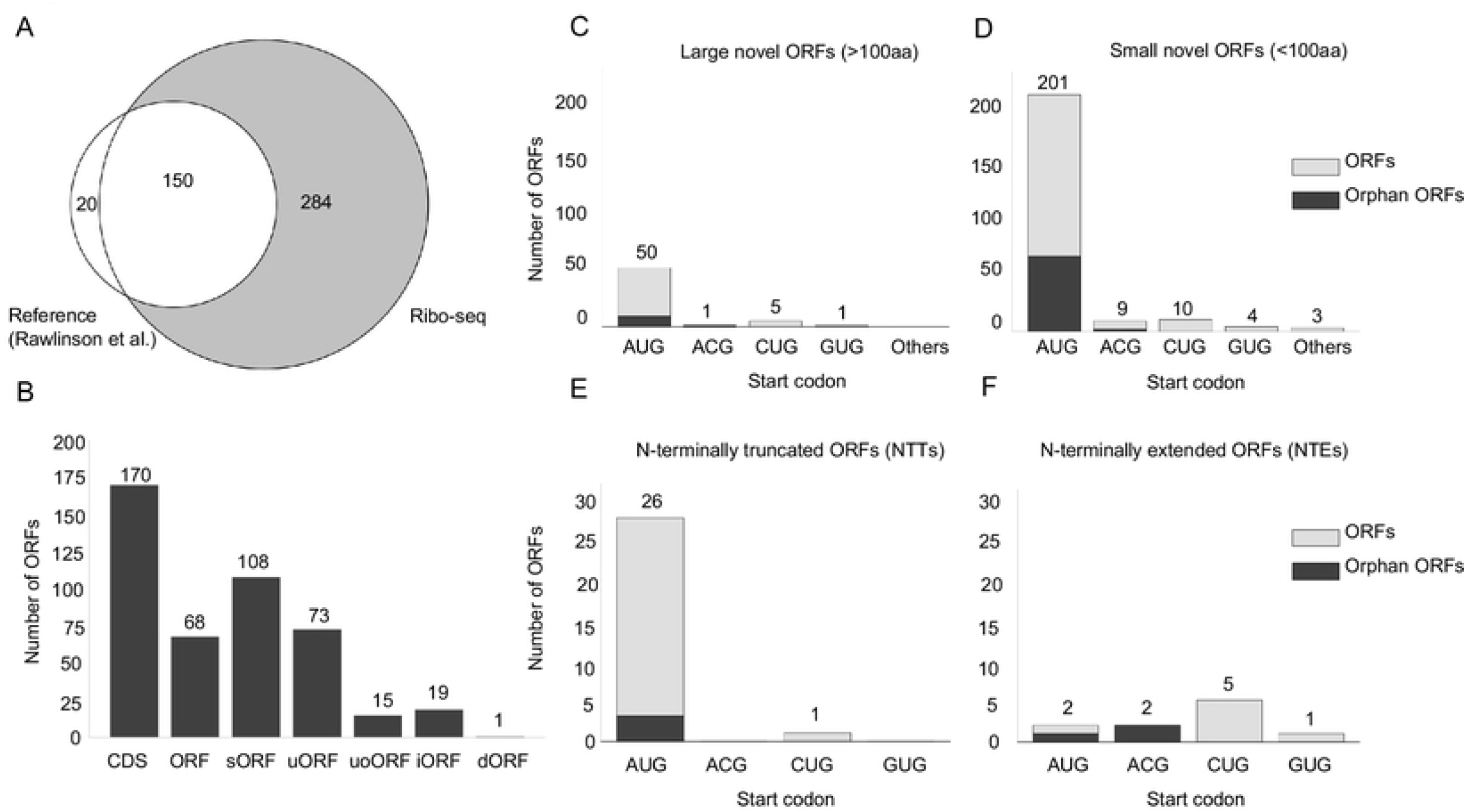
The MCMV translatome. **A.** Venn diagram depicting the number of MCMV ORFs in our revised MCMV genome annotation as detected by ribosome profiling compared to the Rawlinson *et al*. reference annotation [13]. **B.** Total number of viral ORFs annotated by ribosome profiling grouped into CDS (Rawlinson reference annotation), large novel ORFs, N-terminal extensions and truncations (NTEs and NTTs), short ORFs (sORFs), upstream ORFs (uORFs), upstream overlapping ORFs (uoORFs), iORFs (internal ORFs) and downstream ORFs (dORFs). **C-F.** Start codon usage for annotated novel large ORFs, small ORFs, NTEs and NTTs, respectively. ORFs in gray depict orphan ORFs, for which no TiSS could be identified. Each graph depicts the number of ORFs on the y-axis and the start codon usage on the x-axis.

### Characterization of a novel N-terminally truncated ORF in the m145 locus

We identified a novel NTT of the m145 CDS, which we termed m145 ORF #1. This ORF is expressed from a distinct transcript (m145 RNA #1) at 5-fold higher levels than the canonical m145 CDS and lacks the first 340 aa of the 487 aa m145 CDS **(Fig. 7A)**. The glycoprotein encoded by the m145 CDS interferes with NK-cell activation by downregulating the stress-induced NK cell-activating ligand, MULT-I, predominantly in endothelial cells [40]. Considering the immunological significance of this locus, we sought to validate the N-terminally truncated ORF, m145 ORF #1 and assess its role in the regulation of MULT-I. After first validating the m145 proteins through plasmid expression systems **(S7A Fig.)** using V5-tagged ORFs, we generated a C-terminally V5-tagged mutant virus (m145-V5) and analyzed expression in NIH-3T3 and SVEC 4-10 endothelial cells by Western blot **(S7B Fig.)**. This revealed expression of 4 different protein isoforms at ca. 70, 35, 20 and 13 kDa. It is important to note that the canonical m145 CDS encodes a type I membrane protein (55 kDa), which contains a distinct signal peptide and is predicted to undergo N-linked glycosylation [40], thereby explaining the 70 kDa gene product. We then created a panel of virus mutants (**Fig. 7B)** to validate the expression of the two m145 gene products in SVEC 4-10 cells. In particular, mutation of the TATA box of the m145 RNA #1 promoter (Δm145 ORF #1) abrogated the expression of the m145 ORF #1 **(Fig. 7C)**. Interestingly, mutation of this TATA box eliminated all protein isoforms of the m145 locus except for the 70 kDa product encoded by the full-length m145 CDS. On the contrary, the 70 kDa isoform was selectively eliminated when as STOP codon was inserted 40aa downstream of its AUG (Δm145 CDS).

**Fig 7.**
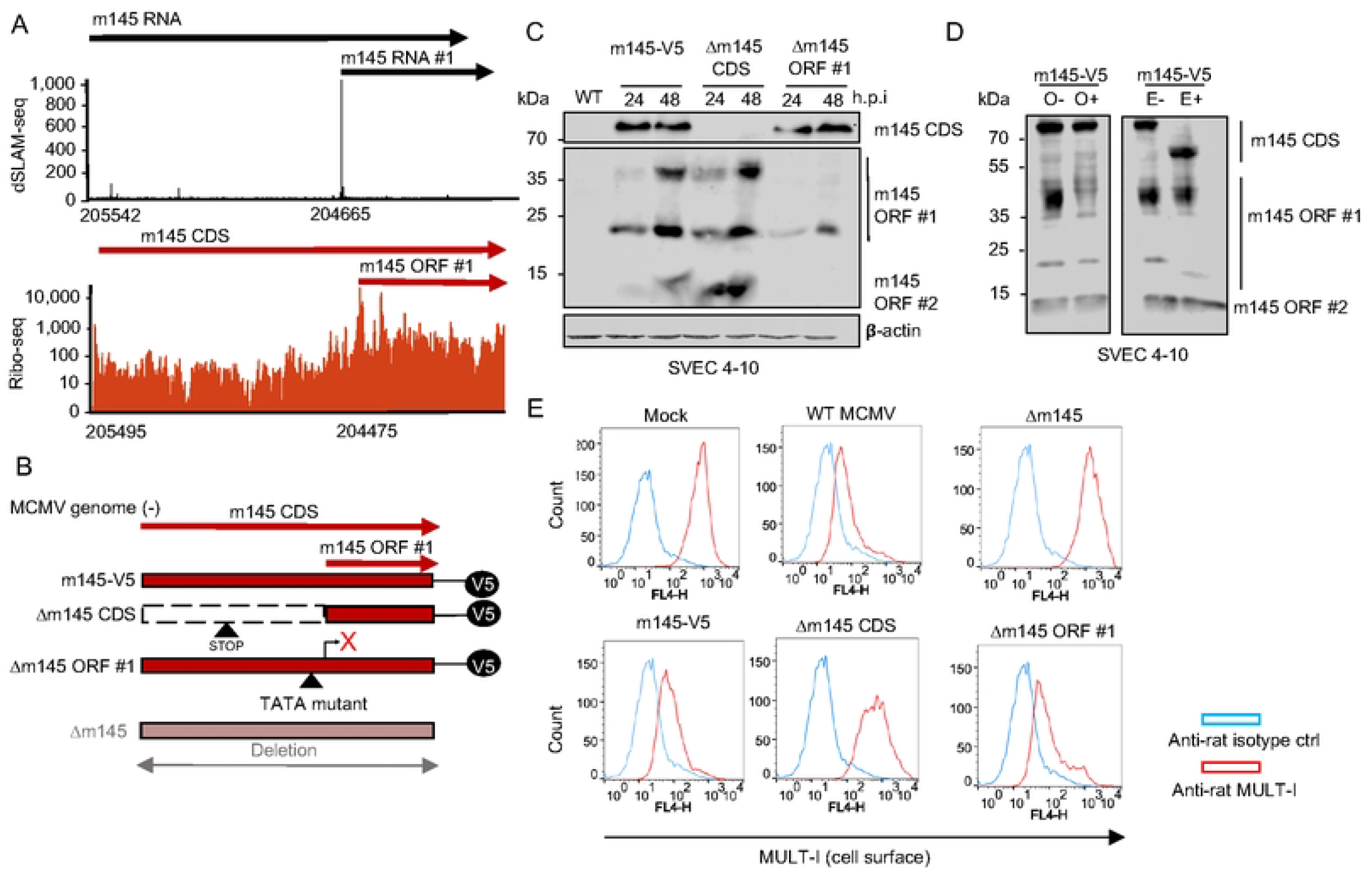
Characterization of N-terminally truncated ORFs in the m145 locus. **A.** ORFs and transcripts expressed from the m145 locus. This includes the so far unknown m145 ORF #1 and #2 expressed from m145 RNA #1. Coordinates for the TiSS and ORF start codon are shown for each transcript and ORF. dSLAM-seq data are shown in linear scale, Ribo-seq data in logarithmic scale. Aggregated reads across all time points mapping to the m145 locus are shown. **B.** Schematic representation of the MCMV mutants generated to characterize novel viral gene products encoded by the m145 locus. Mutant viruses were generated based on a reporter virus with a V5-tag inserted at the C-terminus of the canonical m145 CDS. **C.** SVEC 4-10 murine endothelial cells were infected with the indicated viruses at an MOI of 1 for 24 and 48 h. V5-tagged m145 gene products were characterized by Western blot. Parental WT MCMV infection was used as negative control. **D.** SVEC 4-10 cells were infected for 48 h with the m145-V5 virus at an MOI of 1. Cells were harvested and treated with or without EndoHf (E) or O-glycosidase (O) to qualitatively analyze glycosylation patterns of m145 gene products via Western blot. **E.** SVEC 4-10 cells were infected with m145 virus mutants at an MOI of 1 for 18 h and stained with rat anti-MULT-I and mouse anti-m04 antibodies following cell surface MULT-I analysis through flow cytometry by gating on infected cells (m04+). Anti-rat and anti-mouse isotype antibodies were utilized as negative controls. Western blots and flow cytometry histograms are a representative for two (n=2) and three biological replicates (n=3), respectively.

Next, we asked whether the truncated isoform, m145 ORF #1, predicted to have a molecular weight of 16 kDa and lacking a signal peptide, can undergo glycosylation, thereby explaining the various isoforms. We analyzed glycosylation patterns of the respective proteins through enzymatic treatment with EndoH_f_ and O-glycosidase. This confirmed both the 20 kDa N- and 35 kDa O-linked glycosylated forms of the protein to be encoded by m145 ORF #1 **(Fig. 7D)**. Interestingly, no O-linked glycosylated form of the larger protein encoded by m145 CDS was observed. We hypothesize that its signal peptide marks the protein to exclusively undergo N-linked glycosylation. In contrast, the 13 kDa gene product remained unaffected by glycosidase treatment. While its expression was dependent on the m145 ORF #1 TATA motif and thus translated from m145 RNA #1, mutating the start codon of m145 ORF #1 (m145 ORF #1 mut2) did not disrupt m145 ORF #2 expression **(S7C Fig.)**. We conclude that the 13 kDa gene product results from inefficient ribosome scanning on the m145 RNA #1 and translation initiation at the next AUG start codon located 84 nt downstream. We thus annotated it as an independent small ORF and named it m145 ORF #2 RNA #1 (=m145 ORF #1 translated from RNA #1). These findings indicate the existence of additional truncated viral proteins resulting from variably efficient ribosome scanning.

To clarify which of the m145 ORFs is responsible for downregulation of MULT-I, we analyzed cell surface expression of MULT-I through flow cytometry upon infection with the respective mutant viruses. The Δm145 ORF #1-V5 mutant downregulated MULT-I similar to WT MCMV, indicating that both m145 ORF #1 (despite being expressed at higher levels than m145 CDS) and m145 ORF #2 were not responsible for downregulating cell surface MULT-I and the phenotype was fully attributed to the longer isoform, namely the m145 CDS **(Fig. 7E)**. Our data also confirmed the importance of alternative TiSS usage in governing the expression of MCMV protein isoforms [23].

### Viral uORFs tune viral gene expression

A substantial number of the novel viral ORFs, which we identified via ribosome profiling, represent uORFs, which are located completely upstream of a canonical ORF, and uoORFs, which start upstream and overlap with the canonical ORF. Since translation of u(o)ORFs impacts on translation of their downstream ORFs [41, 42], we aimed to confirm this for selected MCMV u(o)ORFs using dual luciferase reporter assays. We cloned four candidate u(o)ORFs into the psiCheck-2 vector [43] upstream of firefly luciferase. We then mutated their AUG start codon(s) to abrogate translational regulation on the downstream out-of-frame firefly luciferase. This fully relieved translational repression and thus confirmed translation of the m169 uORF (MATp1) [24], m119.3 uORF along with uoORFs in the M35 and M48 locus **(Fig. 8A-D)**. Interestingly, for both the m169 and m119.3 uORF, disruption of the first AUG was not sufficient to fully abrogate their inhibitory potential. However, subsequent mutation of downstream in- and out-of-frame AUG start codons consistently increased downstream luciferase expression. Only when all AUGs (up to 6 for m169 uORF) had been mutated, the observed rescue in luciferase activity matched the expression differences between the respective u(o)ORFs and their larger downstream counterparts observed by ribosome profiling. We conclude that viral u(o)ORFs tune expression of the larger downstream ORFs.

**Fig 8.**
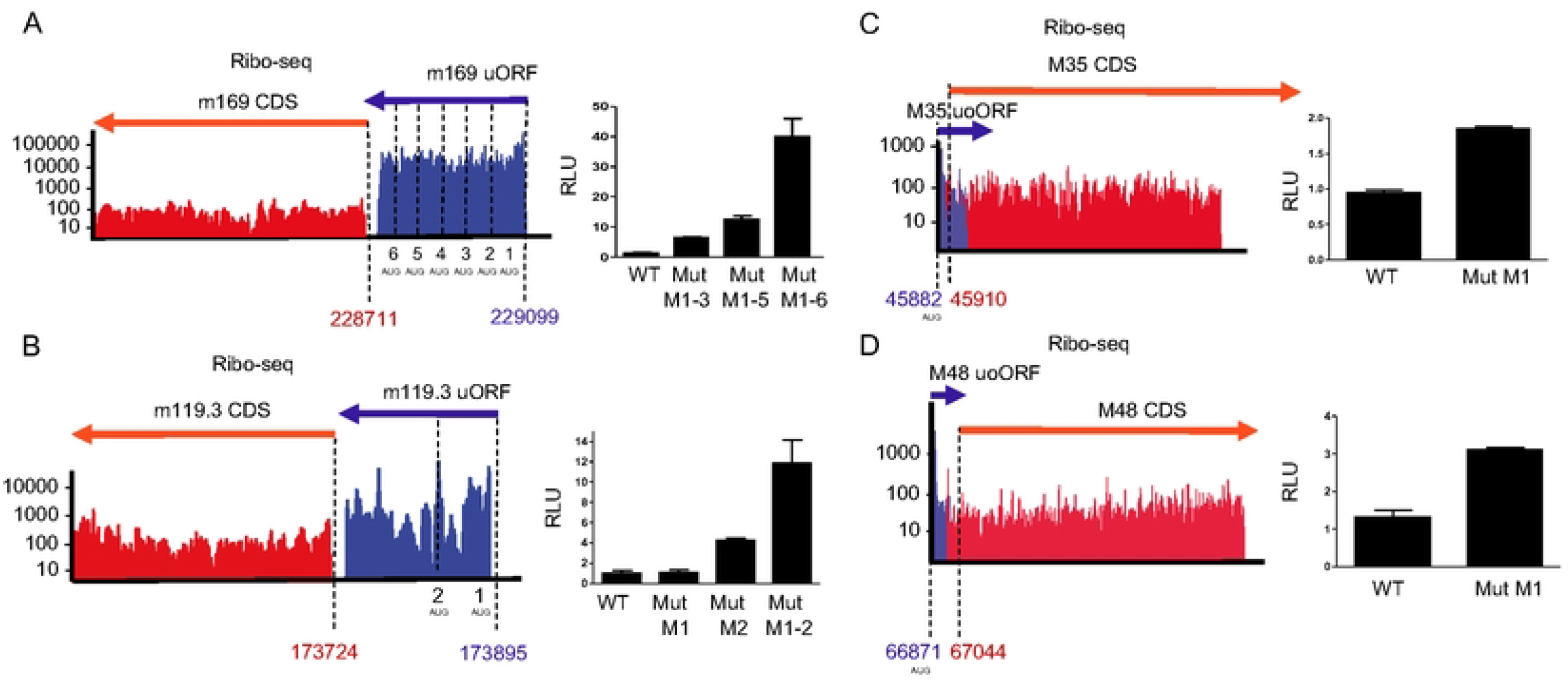
MCMV uORFs/uoORFs tune viral gene expression. Ribo-seq data (aggregated reads in logarithmic scale) of the respective viral genomic loci and their validation by dual luciferase assays are shown for m169 uORF (**A**), m119.3 uORF (**B**), M35 uoORF (**C**) and M48 uoORF (**D**). The number of AUG codons for the respective viral u(o)ORFs are indicated. Coordinates represent the start codons of the u(o)ORFs and ORFs. psiCheck-2 reporter plasmids harbored the indicated MCMV u(o)ORFs (WT) and AUG start codon mutants thereof (Mut) upstream of *firefly-luc* reporter gene. Luciferase assay data at 48h post transfection are shown as mean RLU (Firefly/Renilla ratio) of three biological replicates (n=3) plotted along with the standard error (S.E.M.).

### Reannotation of the MCMV genome

The large number of novel MCMV transcripts and ORFs identified by our approach generated the need for a revised annotation of the MCMV genome. We used the MCMV annotation provided by Rawlinson *et al*. [18] with its 170 viral ORFs as our reference annotation for the BAC-derived pSM3fr MCMV genome sequence [44] curated by our sequencing data. All reference ORF names were maintained accordingly and named as “CDS” (coding sequences) to distinguish these from novel viral ORFs. Any viral ORFs that had previously been revised with minor changes were labelled as “corrected” **(S8 Table)**. We employed the same nomenclature strategy as for the HSV-1 annotation to annotate novel MCMV transcripts and ORFs without altering the existing nomenclature [14]. Briefly, transcription initiating ≥500 nt distant from another transcript was given a new identifier, starting with ‘.5’ to provide room for future additional ORFs in case any TiSS or ORFs had been missed. Transcripts arising from alternative TiSS located within <500 nt upstream or downstream of the main (canonical) transcript in a given locus were labelled as ‘*1’, ‘*2’, … and ‘#1’, ‘#2’, …, respectively. All large novel ORFs were annotated as “ORFs”. Small ORFs were annotated as “uORF”, “uoORF”, “iORF”, “dORF” or “sORF” depending on their relative location to their respective CDS or ORFs. NTEs and NTTs of ORFs were annotated with ‘*’ and ‘#’ respectively. An RNA identifier was used to explain ORFs that could be attributed to alternative TiSS. For example, M25 CDS #1 RNA #1 indicates a truncated ORF (NTT) in the M25 locus translated from an alternative TiSS, namely M25 RNA #1, which initiates downstream of the canonical M25 RNA (**Fig. 2A**). Alternative spliced products were labelled as ORF isoforms (Iso1, Iso2…). ORFs for which no TiSS could be detected were labelled as ‘orphan’. Similarly, transcripts for which no ORF was identified as expressed within the first 500 nt were labelled as ‘orphan’. In total, our final reference annotation includes 66 weakly expressed ‘orphan’ viral RNAs and 88 ‘orphan’ viral ORFs. Reference CDS, which were undetected in our data (and usually lacked a corresponding transcript), were labelled as ‘orphan; not expressed’ but were nevertheless included into the final annotation. The fully reannotated MCMV genome was deposited to the NCBI GenBank Third Party Annotation database.

In summary, promiscuous transcription initiation within the MCMV genome, novel splice isoforms and translation of uORFs and uoORFs upstream of major viral CDS/ORFs explained the vast majority of novel viral gene products identified by our integrative multi-omics approach.

## Discussion

Our study provides a state-of-the-art annotation of the MCMV genome by integrative analyses of a variety of high-throughput sequencing approaches to reveal the hierarchical organization of the entire MCMV transcriptome and translatome at single-nucleotide resolution. While several studies have described novel ORFs and transcripts in previously unannotated regions, our integrative reannotation of the MCMV genome provides a unifying nomenclature for all MCMV gene products. As previously observed for HSV-1, simple peak calling based on our dSLAM-seq and cRNA-seq data would have resulted in the identification of hundreds of additional putative TiSS. While our annotation clearly represents a conservative approach, we restricted the final TiSS to 363 reproducible TiSS by integrative analysis of dSLAM-seq, cRNA-seq and 4sU-seq data. Careful manual inspection of all TiSS candidates in relation to the available Ribo-seq data further increased the reliability of the final TiSS that were included into the new reference annotation. The validity of this approach was confirmed by the strong overrepresentation of Inr elements at the viral TiSS even for the most weakly utilized TiSS. This is consistent with previous findings for HSV-1 and supports the accuracy of our annotation workflow [14]. The vast majority of TiSS were required to explain the expression of novel uORFs, uoORFs, iORFs and splice isoforms, and validated novel NTEs and NTTs revealed by ribosome profiling. Accordingly, only 66 TiSS (of 380, 17.37%) were labelled as orphan while 88 ORFs (of 454, 19.38%) could not be attributed to a viral transcript initiating within 500 nt upstream. We observed a striking number (n=366) of putative splicing events in the MCMV transcriptome. However, the vast majority of these only occurred at relatively low frequencies. We thus decided to include only a conservative 27 splicing events into our new reference annotation.

Clustering transcripts by “new RNA” through dSLAM-seq led to the identification of five distinct clusters describing the kinetics of viral gene expression (Tr0 - Tr4). Clusters Tr0-Tr3 are consistent with the well-described immediate early (Tr0), early (Tr1), early-late (Tr2) and late (Tr3) expression kinetics. Interestingly, dSLAM-seq combined with 4 h of cycloheximide treatment revealed a novel unspliced *ie* gene, namely m166.5 RNA (*ie4*), which we subsequently confirmed by qRT-PCR. The expression of all three *ie* TiSS was significantly enhanced upon inhibition of protein synthesis consistent with a lack of self-inhibition upon CHX treatment. Tr1 and Tr3 promoters were associated with distinct TATA and TATT-box elements, respectively, readily explaining the expression of early and late genes as shown for various herpesviruses. Similar to HCMV [45], the TATT-motif in Tr3 promoters that is recognized by the viral LTF complex tended to be located by about 2 nt further upstream of the TiSS in comparison to the canonical TATA-box motif in the promoters of Tr1 and cellular genes. Tr1 gene expression peaked at 4 hpi following massive repression of transcriptional activity consistent with previous findings [46] describing a similar suppression for MCMV early genes peaking at 3 hpi including the m169 (MAT i.e., Most Abundant Transcript) and m152 genes (>500-fold downregulation) [46]. Transcription of the Tr2 cluster already initiated by 2-4 hpi and thus well before the onset of viral DNA replication at around 12 hpi. However, in contrast to the Tr1 cluster, transcription further increased slightly upon DNA replication and gradually plateaued into late time points. We hypothesize that the presence of both early and late gene expression motifs (TATA-TATT) in Tr2 transcripts explains their consistent expression throughout infection. By mutating the TATA box of an early (Tr1) gene, m152, to a TATT motif, we demonstrate that viral late kinetics and PAA dependence are mediated by the TATT motif and thus the viral LTF complex. However, mutation of a TATA to a TATT motif had little impact on the absolute transcriptional output and did not qualitatively effect transcriptional activity of the m152 promoter early in infection. Other factors, which may include cellular transcription factors activated early in infection, may thus contribute to early viral gene expression.

The absence of a TATT-box element in cluster Tr4 was surprising. The respective transcripts came up significantly later in infection than cluster Tr3 and did not peak until 72 hpi. Their expression is thus unlikely to be dependent on the TATT-specific viral LTF. We hypothesize that transcription initiation of Tr4 transcripts is driven by weak transcription initiation mediated solely by the Inr element in the context of extensive amounts of viral DNA late in infection. Accordingly, even highly expressed Tr4 transcripts were not enriched for any motifs when compared to Tr3 transcripts expressed at similar levels. Studies using conditional LTF knock-out/-down viruses are required to provide further experimental proof for this interesting class of CMV transcripts.

Recently, the Price lab reported on the identification of ≈7,500 transcription start site regions (TSRs) in the HCMV genome during lytic infection of fibroblasts, which corresponds, on average, to a TSR every 65 nt [47], using PRO-Cap-seq. These were corroborated by additional studies attributing their expression kinetics at least in parts to the viral IE2 protein and LTF [10, 48]. While our TiSS profiling data do not exclude the presence of a much larger set of TSRs for MCMV, the TiSS we identified (***i***) correspond to stable RNAs and (***ii***) are largely sufficient to explain the complete MCMV translatome identified by ribosome profiling. Furthermore, the presence of thousands of additional stable viral transcripts should have resulted in the translation of hundreds of additional viral ORFs observable by Ribo-seq. We conclude that the number of stable MCMV transcripts that are actively translated is unlikely to exceed our annotation by an order of magnitude. Importantly, the PRO-Cap approach not only detects stable transcripts but also transcription of highly unstable transcripts including promoter- and enhancer-derived RNAs. Interestingly, dRNA-seq and STRIPE-seq analysis on HCMV-infected fibroblasts, which both only detect stable transcripts, only confirmed ≈1,700 of the >7,000 TSRs but indicated extensive non-productive (pervasive) transcription of the HCMV genome [45]. Our data for MCMV are consistent with our findings for HCMV showing that a large fraction of the >7,000 TSRs reported for HCMV presumably do not correspond to stable viral transcripts. It will be interesting to study whether transcription initiation is as promiscuous in lytic MCMV infection as observed for HCMV.

Our TiSS profiling data provide strong additional evidence for the newly identified ORFs and small ORFs detected by Ribo-seq. In the vast majority of cases, the respective novel ORFs initiate from the first AUG downstream of the respective TiSS. An excellent example of this is m145 ORF #1. It is translated from a so far unknown viral transcript (m145 RNA #1) that initiates in the middle of the m145 CDS. However, as we demonstrated for the 13 kDa m145 ORF #2, inefficient ribosomal scanning of m145 RNA #1 may also result in translation initiation at the next downstream AUG resulting in the expression of a truncated protein isoform. Although the less abundantly expressed m145 CDS was responsible for the published effects on MULT-I [40], our findings confirm expression of at least two additional viral proteins (m145 ORF #1 and #2) and implicate differentially glycosylated gene products expressed from the m145 locus. While we were a bit surprised to see that the less prominently expressed m145 CDS accounted for the reported regulation of MULT-I, high expression of m145 ORF #1 may well have confounded the interpretations of previous *in vivo* experiments [49]. Further studies are required to functionally characterize the role of the additional proteins expressed from the m145 locus.

Similar to HCMV [13], 227 of 284 novel MCMV ORFs (80%) were <100 aa in size, a substantial fraction of which represented uORFs or uoORFs. Their cellular counterparts have been implicated to control gene expression of their downstream ORFs at the translational level [41, 42]. By identifying both the u(o)ORFs and their corresponding TiSS, our data will now enable functional studies pertaining to u(o)ORF-mediated gene regulation in CMV infection. Although many of the novel ORFs that we identified are less than 100 aa in size, they may nevertheless encode for abundant microproteins with important functions. The potential of such novel sORF-encoded viral microproteins for productive infection was recently demonstrated for the m169 uORF encoding an NK cell immune evasin [24] and the m41.1 gene product [50] that blocks mitochondrial apoptosis. Mass spectrometry and structural biology data should thus be reanalyzed to look for novel CMV microproteins. Finally, small ORFs have also been implicated to generate antigenic peptides, resembling rapidly generated DRiP-derived peptides [51]. Such peptides generated from microproteins may form a major component of the antigenic repertoire [38, 51, 52], playing a role in various diseases [17]. Our revised annotation of the MCMV genome now enables to assess their role in antigen presentation and immune evasion in the MCMV model.

## Materials and Methods

### Cell culture, viruses and infection

NIH-3T3 (ATCC® CRL-1658™) Swiss mouse embryonic fibroblasts were grown in DMEM (Dulbecco’s Modified Eagle’s Medium) supplemented with 100 IU/mL penicillin (pen), 100 μg/mL streptomycin (strep) and 10% NCS (New-born calf serum). M2-10B4 (ATCC® CRL-1972™) fibroblasts were grown in RPMI-1640 (Roswell Park Memorial Institute Medium) supplemented with 100 IU/mL pen, 100 μg/mL strep and 10% FCS (Fetal calf serum). 293T (ATCC® CRL-3216™) human embryonic kidney (HEK) epithelial cells and SVEC 4-10 mouse endothelial cells (ATCC® CRL-2181™) were grown in DMEM supplemented with 100 IU/mL pen, 100 μg/mL strep and 10% FCS. All cells were grown in 5% CO_2_ at 37°C. All viruses were generated by infecting M2-10B4 cells after virus reconstitution. BAC-derived MCMV Smith strain was utilized for all sequencing experiments [44]. Infected cells and supernatants were harvested after >90 % infection for virus purification and titration of virus stocks was conducted by standard plaque assays on NIH-3T3 cells [3]. The Δm145 virus has been published previously [49]. Infections were conducted using centrifugal enhancement at 800g for 30 min in 6-well plates followed by incubation at 37°C in 5% CO_2_ for 30 min. Media change following incubation marked the 0-hour time point of infection. An MOI of 10 was used for all high-throughput experiments.

### Virus mutagenesis and reconstitution

The MCMV Smith strain bacterial artificial chromosome (BAC) in GS1783 *E. coli* [44] was used to construct MCMV virus mutants using en passant mutagenesis [53], as described previously. Selected clones were verified by restriction enzyme digestion and Sanger sequencing of the respective locus. BAC DNA was purified using the NucleoBond BAC 100 kit (Macherey-Nagel #740579) and were transfected into early passage NIH-3T3 cells in 6 well plates using TransIT-X2® dynamic delivery transfection system (Mirus). Viruses from cell culture supernatants were passaged on M2-10B4 cells followed by virus purification and titration [3]. All primers along with cloning strategy utilized are described in **S9 Table**.

### RT-qPCR analysis

Wild-type MCMV infections were performed as described for dSLAM-seq in 12-well plates using centrifugal enhancement at 800g/30 minutes. Cycloheximide (50 μg/mL) treatment was performed at 0hpi. DMSO was used as mock treatment. Samples were harvested at 4hpi, followed by RNA extraction using the Zymo Quick™ Microprep kit including an additional gDNA digestion step using TURBO™ DNAse (Life technologies). 300-400 ng RNA was used to prepare cDNA utilizing the Bimake 5X qRT All-in-one- cDNA synthesis mix. A 1:5 dilution of the obtained cDNA was subject to 2-step qPCR using the SYBR green qPCR MasterMix (2X) by MedChemExpress as described by the manufacturer. qPCR was performed on the Roche LightCycler® 96. Each qPCR included two technical replicates per gene. The obtained data were analyzed by ddCt analysis for three biological replicates. Mean and SEM were plotted using Graphpad Prism. Primers used are listed in **S9 table**.

### Plasmids and transfection

The psiCheck-2 vector was utilized for validating uORFs/uoORFs by dual luciferase assays [43]. All uORF/uoORF constructs were purchased as gene block fragments from Integrated DNA Technologies (IDT) bearing homologies to psiCheck-2 BstBI and ApaI sites. Cloning was performed using the In-fusion® HD Cloning Plus kit (Takara Bio) as per manufacturer’s instructions, followed by transformation in Stellar competent cells (Takara Bio). uORF/uoORF start codon mutants were generated by double-fragment infusion cloning using two PCR products bearing homologous ends containing mutations. MCMV m145 ORFs were cloned into pCREL-IRES-Neon expression plasmids with a C-terminal V5-tag between Spe-I and Cla-I restriction sites using infusion cloning. All plasmids were sequenced and purified using the PureYield™ Promega Midiprep system. For luciferase assays, plasmids were transfected in NIH-3T3 cells in a 96-well plate using Lipofectamine ™ 3000 (Invitrogen). Luciferase readings were measured 48 hours’ post-transfection using the Dual-Glo® Luciferase assay system (Promega), as per manufacturer’s instructions using the Centro XS^3^ LB960 system (Berthold Technologies). For Western blot, 6-well plates seeded with HEK293T cells were transfected with the m145 containing expression plasmids using TransIT-X2® dynamic delivery transfection system (Mirus) and cells were harvested at 48 hours’ post transfection. All primers and synthetic constructs used are described in **S9 Table**. All restriction enzymes were purchased from NEB. Luciferase data mean values (Firefly/Renilla ratio) were plotted along with standard error (S.E.M) as relative light units (RLU) for three biological replicates using Graphpad Prism.

### Western blot

Cells were lysed with 2X Laemmli sample buffer (Cold Spring Harbour protocols) with 20% β-Mercaptoethanol. Lysed samples were sonicated and heated at 95°C/10 minutes. Tris-Glycine SDS-PAGE (12%) and wet transfer (Tris-Glycine-20% Methanol) on 0.2 μm Nitrocellulose membrane (Amersham™ Protran™) were performed using the Mini Gel Tank (Life technologies). Membranes were subsequently subject to blocking in 5% (v/v) skimmed milk in 1X PBST (Phosphate buffered saline – 0.1% Tween 20) at room temperature for one hour. Samples were probed with rabbit anti-V5 antibody (Cell Signalling #13202S) at a 1:1000 dilution, overnight at 4°C and then probed with a 1:1000 dilution of α anti-rabbit IgG-Horseradish peroxidase (HRP) – Sigma Aldrich A0545. All antibodies were diluted in 5% (v/v) milk in 1X PBST. Proteins were analyzed by visualizing the blots on LI-COR Odyssey® FC Imaging System. For O-glycosidase (NEB P0733S) treatment, samples were lysed in 1X RIPA lysis buffer containing anti-protease cocktail (cOmplete™, Mini Protease Inhibitor Cocktail, Roche) along with denaturing buffer supplied by NEB. Treatment with O-Glycosidase and Neuraminidase (NEB P0720S) was conducted as per manufacturer’s instructions for one hour at 37°C. A similar protocol was performed for EndoH_f_ (NEB P0703S). ß-actin was used as a housekeeping control and immunoblotting was performed using mouse anti-ß-actin primary monoclonal antibody (C4-sc-47778 Santa Cruz Biotechnology, Inc.), and the fluorescent IRDye® 680 RD goat anti-mouse IgG (Licor) was used as a secondary antibody. Both antibodies were diluted 1:1000 in 1X PBST. All western blot images were processed through ImageStudio Lite.

### Flow cytometry

Uninfected and MCMV-infected SVEC 4-10 were washed with 1X PBS and detached using TrypLE™ Express (Gibco) 18 hpi followed by blocking in 10% FCS-PBS (1X) for 30 minutes. Cells were stained with rat anti-MULT-I and/or mouse anti-MCMV m04 at a dilution of 1:100 including isotype controls for MULT-I (eBioscience™ Rat IgG2a kappa control eBR2a) and m04 (eBioscience™ Mouse IgG2b kappa control eBMG2b) as well as only secondary antibody controls by incubating for 30 minutes on ice. Both anti-MULT-I and anti-m04 antibodies were provided by Stipan Jonjic. Followed by primary antibody staining, cells were stained by Invitrogen Goat anti-Rat IgG (H+L) Alexa Fluor 647 (MULT-I) and/or Abcam goat polyclonal anti-Mouse Alexa Fluor 488 (m04) at a dilution of 1:1000 for 30 minutes on ice. All antibodies were diluted in 10% FCS-PBS (1X). Cells were finally suspended in FACS buffer (1X PBS with 0.5% BSA, 0.02% sodium azide). Flow cytometry was performed using the BD Biosciences FACS Calibur™ Cell Quest Pro system. Gating and further analysis was performed using FlowJo™10. Briefly, live SVEC 4-10 cells were gated for anti-mouse Alexa Fluor 488 bound MCMV infected cells via the FL-1 channel (488 nm Argon ion laser and 530/30 filter) followed by histogram visualization of cell surface expression levels of MULT-I bound by anti-rat Alexa Fluor 647 using the FL-4 channel (635 nm Red diode laser and 661/16 filter). Flow cytometry analysis was similarly performed by analyzing GFP (FL-1) and mCherry expression (FL-3), post fixing in 4% formaldehyde and MFI values and S.D. for each time point/condition were plotted using Graphpad Prism for three biological replicates. Prior to fixing, the samples were analyzed qualitatively via microscopy at 10X resolution using the Leica DMi8 system.

### Transcription start site (TiSS) profiling

Cycloheximide treatment at 50 μg/mL was conducted at the time of infection and phosphonoacetic acid (PAA) treatment was conducted at 300 μg/mL one-hour post infection. cRNA-seq and dSLAM-seq were performed as described [14] with minor modifications. For all dSLAM-seq samples, 4sU labelling was initiated by adding 400 μM for 60 minutes before harvest using TRI reagent (Sigma Aldrich) as described by manufacturer and purified by standard phenol-chloroform extraction. Total RNA was re-suspended in 1X PBS buffer. U-to-C conversion were initiated by iodoacetamide (IAA) treatment as described previously [28] and RNA was re-purified using RNeasy MinElute (Qiagen). Efficiency of IAA conversion was checked by converting 1mM 4sU and analyzing the change in absorption (loss of adsorption maximum at 365 nm) upon IAA treatment [28]. Following this, library preparation using the dRNA-seq protocol and Xrn-I digestion was performed by the Core Unit Systems Medicine (Würzburg) as described previously for HSV-1 [14]. Sequencing was performed on NextSeq500 (Illumina). For cRNA-Seq, the same protocol was utilized as for HSV-1 [14]. 5’ read enrichment was obtained using chemical RNA fragmentation (50-80 nt fragments) and libraries were prepared using 3’ adaptor ligation and circularization. Libraries were sequenced on a HiSeq 2000 at the Beijing Genomics Institute in Hong Kong. Total RNA-seq and 4sU-seq was conducted as described[26]. Briefly, 4sU labelling was conducted at 500μM for 60 minutes for the time points described in **Fig. 1**. Cells were lysed in Trizol (Invitrogen) and total and 4sU labelled (newly transcribed RNA) were isolated as per previous protocols. Libraries were prepared using the stranded TruSeq RNA-Seq protocol (Illumina, San Diego, USA) as described and libraries were sequenced by synthesis sequencing at 2 × 101 nt on a HiSeq 2000 (Illumina).

### Ribosome profiling

Ribosome profiling time-course (lysis in presence of cycloheximide) experiments were conducted as described [13] for time-points as shown in **Fig. 1** for four biological replicates. Additionally, translation start site (TaSS) profiling was performed by culturing cells in medium containing either Harringtonine (2 μg/ml) or Lactimidomycin (50 μM) for 30 min prior to harvesting. Two biological replicates were generated for Harringtonine pre-treatment and one for Lactimidomycin. Libraries were generated as described for cRNA-Seq [14], which introduces a 2 + 3 nt unique molecular identifier (UMI), facilitating the removal of PCR duplicates from sequencing libraries. All libraries were sequenced on a HiSeq 2000 at the Beijing Genomics Institute in Hong Kong.

### Data analysis and statistics

Random and sample barcodes in cRNA-seq and ribosome profiling data were analyzed by trimming the sample and UMI barcodes and 3’ adapters from the reads using our in-house computational genomics framework gedi (available at https://github.com/erhard-lab/gedi). Barcodes introduced by the reverse transcription primers included three random bases (UMI part 1) followed by four bases of sample-specific barcode followed by two random bases (UMI part 2). Reads were mapped using bowtie 1.2 against the mouse genome (mm10), the mouse transcriptome (Ensembl 90), and MCMV (KY348373, checked and corrected according to mutations listed in the previous publication [44]). Reads were assigned to their specific samples based on the sample barcode. Barcodes not matching any sample-specific sequence were removed. PCR duplicates of reads mapped to the same genomic location and sharing the same UMI were collapsed to a single copy. Two observed UMIs that differed by only a single base are likely due to a sequencing error and were therefore considered to be the same UMI. If the reads at this location mapped to k locations (i.e., multi-mapping reads for k > 1), a fractional UMI count of 1/k was used.

dRNA-SLAM-seq and 4sU-seq data were processed similar to cRNA-seq and ribosome profiling data with the exception of STAR (v.2.5.3a) being used to map the reads and PCR duplicates were not collapsed as no UMIs were used.

Our dRNA-SLAM-seq and cRNA-seq TiSS profiling data were analyzed with our TiSS analysis pipeline iTiSS (available at https://github.com/erhard-lab/iTiSS) [30], which identifies potential TiSS at single-nucleotide resolution. The SPARSE_PEAK module was used for dRNA-SLAM-seq. For cRNA-seq data, DENSE_PEAK, DENSITY, and KINETIC modules were used. For each replicate, reads were pooled from all time points. Subsequently, for each dataset, TiSSMerger2, a subprogram in iTiSS, was used to merge TiSS with a +/- 10 bp window. Correspondingly, all TiSS from all datasets were merged using TiSSMerger2 also with a +/- 10 bp window. iTiSS assigned a score ranged from 1 to 4 for each TiSS based on several criteria:

i. Significant accumulation of the 5’-end of reads in both replicates of the dRNA-SLAM-seq dataset at the TiSS (SPARSE_PEAK module).
ii. Significant accumulation of the 5’-end of reads in both replicates of the cRNA-seq dataset at the TiSS (DENSE_PEAK module).
iii. Stronger transcriptional activity downstream than upstream of the potential TiSS in both cRNA-seq replicates (DENSITY module).
iv. Significant temporal changes in TiSS read levels during the course of infection in both cRNA-seq replicates (KINETIC module). We also add 3 more criteria for scoring.
v. Stronger transcriptional activity downstream than upstream of the potential TiSS in both 4sU-seq replicates.
vi. Significant temporal changes in TiSS read levels during the course of infection in both 4sU-seq replicates.
vii. The presence of an ORF at most 250 bp downstream, which was not yet explained by another transcript.

Thus, in total, we assigned a score between 1 to 7 for each TiSS. We then manually inspected the final list of TiSS using MCMV genome browser and selected TiSS with a prominent signal to be included in our annotation. A histogram was created showing the number of criteria fulfilled by all annotated TiSS. In addition, we also created a heat map and a bar plot to compare cRNA-seq and dRNA-SLAM-seq by calculating the enrichment of reads at TiSS compared to +/- 100bp region around the TiSS (S1A Fig). Both figures indicate that dRNA-SLAM-seq provides a better signal-to-noise ratio compared to cRNA-seq.

The total RNA count for each annotated TSS was calculated by counting the number of reads whose 5’ end is inside a +/- 5 bp window of a given TiSS. Subsequently, for each transcript, Uridine to cytosine (U-to-C) conversion rates, error rates, and new to total RNA rates (NTRs) were estimated by analyzing dRNA-SLAM-seq data using GRAND-SLAM [29]. Only reads with 5’ ends inside +/- 5 bp window of an annotated TiSS were considered. Newly synthesized RNA count of each TiSS was then calculated by multiplying NTR value with total RNA count obtained from dRNA-SLAM-seq data.

We grouped TiSS into three groups based on their expression level. For each group, we generated sequence logos from the −34 to +5 window around TiSS using WebLogo [54].

All n=363 TiSS were clustered using the k-means clustering algorithm [55] into four transcription classes (Tr) based on the new RNA expression. Clustering was repeated 10,000 times with different random initializations. Cluster centroids from each clustering replication were then clustered again one more time to obtain a consensus centroid. This consensus centroid was used for the final clustering of TiSS.

For each TiSS cluster, promoter motifs and their location were searched using MEME [56] with *-evt 0.05, -nmotifs 5, -minw 6*, and *-maxw 15* parameters. We searched the motifs inside the −34 to +5 window of a given TiSS. In addition, we also generated sequence logos from each cluster using the same window. Tr3 and Tr4 were analyzed further by grouping each of them into four groups based on the expression value. Sequence logos were generated using the same procedure as mentioned before.

We used our in-house tool PRICE version 1.0.4 [38]to predict MCMV ORFs. A list of putative ORFs was then manually inspected by using MCMV genome viewer to select *bona-fide* ORFs which will be included in the final annotation. We grouped these ORFs into CDS (ORFs which are included in previous annotation), ORF (ORFs with length ≧ 100 amino acids (aa) which are not in previous annotation), sORF (ORFs with length < 100 aa), uORF (ORFs located upstream of the canonical ORF, but inside the transcript region), uoORF (ORFs located upstream of the canonical ORF and also overlap the canonical ORF but in a different frame), iORF (ORFs located inside a canonical ORF but in a different frame), and dORF (ORFs located downstream of the canonical ORF, but inside the transcript region).

### Identification of poly(A) sites and splicing events

4sU-seq reads were first filtered for rRNA reads by aligning reads against rRNA sequences using BWA [57] with a seed size (parameter -k) of 25. If both reads in a read pair aligned to rRNA without errors, they were removed from further analysis. Filtered 4sU-seq reads and all total RNA-seq reads were aligned against the MCMV genome using ContextMap version 2.7.9 [58] (using BWA as short read aligner and allowing at most 5 mismatches and a maximum indel size of 3). ContextMap identifies also reads containing to part of the poly(A) tail and predicts poly(A) sites from these reads as previously described [58]. Default parameters were used for poly(A) site prediction. Candidate splice junctions were predicted if >10 reads were identified by ContextMap in at least one sample that overlapped at least 10 nt on both sides of junction. All viral introns are listed in **S3 Table.** All viral poly(A) sites are listed in **S10 Table.**

## Code availability

The gedi toolkit, which was used for mapping and most of the analysis steps, is available on GitHub (https://github.com/erhard-lab/gedi). iTiSS, which is a module for gedi is available separately on GitHub (https://github.com/erhard-lab/iTiSS). The source code of all additional custom scripts generated for generating the Figures, tables and analyzing the data in general can be found at Zenodo (https://doi.org/10.5281/zenodo.6861955). A genome browser including all data is available at https://doi.org/10.5281/zenodo.7105431.

## Data availability

All sequencing data produced in this study are available at GEO (accession number GSE212289).

## Acknowledgements

This work was supported by a grant from the Deutsche Forschungsgemeinschaft (FOR 2830, DO 1275/7-1 and ER 927/1-1 to LD and FE, respectively) and grant FR2938/9-1 to CCF. The funders had no role in study design, data collection and analysis, decision to publish, or preparation of the manuscript.

## Author contributions

Conceptualization, investigation and development of methodology- LD, FE, ML, IM, CJ, AR, AH, SJ. Data curation and formal analysis- LD, FE, ML, IM, CJ, CCF, VJL. Validation- ML, IM, BKP, TH, AM, AG. Visualization- ML, IM, FE and LD. Writing – ML, IM, LD, FE. Funding acquisition- LD and FE. Supervision- LD and FE.

## Supporting information

### Supporting Files

**S1 File: Description of high-throughput sequencing datasets used in this study.**

### Supporting Tables

**S1 Table: List of all MCMV transcripts.** List of all identified and annotated viral transcripts. TSS = transcription start site; TTS = transcription termination site.

**S2 Table: List of all splicing events annotated and identified through 4sU-seq analysis.** List of all splicing events that were included into the new MCMV reference genome annotation.

**S3 Table: List of putative introns detected by 4sU-seq. List of all putative splicing events that were not included into the new MCMV genome reference annotation.**

**S4 Table: Immediate-early viral transcripts.** Enrichment of immediate early transcripts upon cycloheximide (CHX) treatment. Immediate-early (*ie*) transcripts were determined by analyzing nascent RNA with and without cycloheximide pre-treatment. *ie* Transcripts unaffected by cycloheximide are highlighted in red.

**S5 Table: List of all MCMV ORFs.** List of all MCMV ORFs including their name, ORF type, coordinates, strand and length of predicted protein products.

**S6 Table: List of all unidentified CDS predicted by Rawlinson *et al*.** List of all CDS predicted by Rawlinson et al. that were not observed in our data. Their respective CDS were nevertheless included in our revised MCMV genome annotation as ‘not expressed/orphan’ CDS. * CDS that are overlapping other MCMV genes by greater than 60% and are thus less likely to be protein coding and have no homologs in herpesviruses or cellular proteins as described by Rawlinson et al.

**S7 Table: List of previously unannotated ORFs confirmed by us and validated in several studies**. Table of all MCMV ORFs that have been identified by others and that were confirmed by our data.

**S8 Table: Detected ORFs with minor corrections i.e. ‘CDS (corrected)’ as verified by previous studies.** Table of all previously reported MCMV CDS that required minor corrections based on our data. * Initiation at a downstream AUG

**S9 Table: List of primers and gene constructs used.**

**S10 File: List of MCMV poly(A) sites.** List of all MCMV poly(A) sites that were annotated based on untemplated adenines on sequencing reads.

### Supporting Figures

**S1 Fig: Characterization of the MCMV transcriptome.**

**A.** Heat maps comparing read enrichment at transcription start sites (TiSS) in the cRNA-seq and dSLAM-seq data. **B**. Histogram depiction of the number of MCMV TiSS satisfying the indicated number of criteria of the iTiSS algorithm. A detailed description of the employed criteria is included in methods.

**S2 Fig: Examples of MCMV splicing events.**

Each schematic depicts viral gene expression and splicing in a given locus. Aggregated reads of Ribo-seq, cRNA-seq and dSLAM-seq data across all time points of infection are shown. Ribo-seq data are indicated in logarithmic scale, cRNA-seq and dSLAM-seq data in linear scale. The arrows at the top depict the annotated of transcripts (black) and ORFs (colored depending on the translated frame (yellow, purple and green)). The bold dotted line represents the introns detected by 4sU-seq. **A**. In the m133 locus, splicing of two introns leads to the expression of both a known (Iso1) and a novel spliced ORF (Iso2), the latter is expressed through an alternative donor site, as predicted by Rawlinson *et al*. Both transcripts are expressed with early kinetics **B**. In the M116 locus, splicing explains a truncated M116 CDS Iso2 (M116.1p) revealed by ribosome profiling, whose transcript may terminate at an earlier poly A site (M116 RNA Iso2). A polyA site downstream explains translation of the unspliced ORF. Here, transcription continues past the 1^st^ poly A site (PAS) resulting in M116 RNA Iso1. **C**. In the m147.5 locus, splicing leads to the expression of a previously validated spliced ORF. **D**. In the m124 locus, splicing necessitates correction of the previously annotated m124 ORF. Coordinates of the start codons and splicing acceptor and donor sites are displayed.

**S3 Fig: Splicing events in the m60-m73.5 locus.**

Graphs represent TiSS profiling data (black) from cRNA-seq and dSLAM-seq as well as ORFs called by Ribo-Seq (different colors represent different frames of translation). Aggregated reads of Ribo-seq, cRNA-seq and dSLAM-seq data across all time points of infection are shown. Ribo-seq data are indicated in logarithmic scale, cRNA-seq and dSLAM-seq data in both linear and logarithmic scale. Spliced ORFs are depicted by exons connected with a dotted line representing introns at the bottom. Multiple splicing events were observed in the m60-73.5 locus, of which the m60 RNA and M73-m73.5 spliced transcripts have already been validated previously (see **S2 Table**). Of note, translation occurs in different fames upstream of splicing thereby explaining translation in different frames in the common downstream exon. For a given frame of translation at the second exon, the expression levels correlated well with the respective upstream exons.

**S4 Fig: The m166.5 RNA constitutes a novel *ie* gene (*ie4*).**

**A.** Cycloheximide treatment combined with dSLAM-seq identified a so far unknown viral immediate early transcript in the m166.5 locus. Aggregated reads of Ribo-seq, cRNA-seq and dSLAM-seq data across all time points of infection are shown. Ribo-seq data are indicated in log scale, cRNA-seq and dSLAM-seq data (-/+CHX) in linear scale. The m166.5 immediate-early transcript (ie4) and its corresponding m166.5 ORF overlap with the m167 CDS (orphan) and partially overlap with the N-terminal part of the m166 CDS. **C.** Line graphs representing gene expression (new RNA levels) of *ie* genes over time for two replicates per gene.

**S5 Fig.: Characterization of the Tr4 cluster of viral transcripts.**

**A.** Quantile groups segregated according to levels of expression for Tr3 and Tr4 transcripts on the basis of new RNA for the respective viral TiSS obtained from the dSLAM-seq data. The x-axis displays hours post infection. Relative expression is shown on the y-axis. Tr3 and Tr4 transcripts are indicated by green and blue lines, respectively. **B.** Motif analysis (MEME) for all four quantiles for Tr3 and Tr4 transcripts. **C.** Comparison of MEME Motif analysis for the top thirty transcripts of Tr4 compared with Tr3 transcripts of corresponding expression levels confirms the lack of a TATA box-like motif.

**S6 Fig.: Gene expression in the viral *ie2* locus.**

**A.** Schematic of the *ie2* locus. The canonical TiSS is represented by the dominant spliced *ie2* transcript (m126-m128 RNA) comprising 2 introns and only one *ie2* coding exon initiating at the first AUG (186087) shown i.e., m128 CDS (*ie2* Exon 3). A second AUG represents a truncated isoform (m128 CDS #1 RNA #1). Aggregated reads of Ribo-seq, cRNA-seq and dSLAM-seq data across all time points of infection are shown. Ribo-seq data are indicated in log scale, cRNA-seq and dSLAM-seq data in linear scale. **B.** Graphs represent TiSS profiling data (black) from dSLAM-seq including kinetics for 6 and 24 hpi and ORFs called by Ribo-seq (Colored) for the same time points for a given replicate. The arrows above depict manual annotations of transcripts and ORFs. Alternative transcription initiation at the *ie2* (m128) locus led to the expression of an N-terminally truncated ORF expressed from an early TiSS (m128 RNA #1) whose expression was not influenced by PAA treatment. cRNA-seq and dSLAM-seq data are represented in linear scale, Ribo-seq in logarithmic scale.

**S7 Fig: Validation of m145 ORFs**

Both the m145 CDS and m145 ORF #1 were cloned into expression plasmids (pCREL-IRES-Neon) with their expression driven by a CMV promoter. **A.** Expression of the two viral ORFs was validated via transfection of the respective plasmids into HEK293T cells. Western blots were performed at 48 h post transfection. **B.** The m145-V5 virus described in **Fig 5B** was used to infect both NIH-3T3 and SVEC 4-10 cells at an MOI of 1 for the respective time points to validate the m145 gene products, whose expression was similar in both cell lines. **C**. A start codon mutant of m145 ORF #1 (Δm145 ORF #1 mut 2) was utilized to infect SVEC 4-10 cells for 48 h. Western blot analysis revealed expression of m145 ORF #2 to remain unaffected. WT indicates wild-type MCMV. β-actin was used as a housekeeping control. Images are a single representative of 2 biological replicates (n=2) for each experiment.

